# Composing the value signal for dopamine-mediated learning

**DOI:** 10.1101/2025.10.10.681616

**Authors:** Pranav Mahajan, Ben Seymour

## Abstract

The seminal reward prediction error account of dopamine has been highly successful, but faces several key challenges. Most notable are the difficulty of learning multiple rewards simultaneously, inefficient on-policy learning, and accounting for the heterogeneous striatal responses observed across and within striatal targets. Here we address these issues with a normative entropy-regularised reinforcement-learning framework. We propose that dopamine optimises not just cumulative rewards, but a reward value function augmented by a penalty for deviating from a default behavioural policy. In simulations, this off-policy formulation provides a principled solution to composing multiple reward values, avoids the interference and unintended unlearning seen in multi-objective on-policy methods when priorities change, and adapts more efficiently than standard alternatives in environments with non-stationary rewards. More broadly, the framework offers a unified account of dopamine heterogeneity between and within striatal targets, including a normative way to understand why aversive and action prediction errors may coexist in the tail of the striatum. Together, these results suggest that dopamine-mediated learning may be better captured by prediction errors in composable, entropy-regularised value functions than by a single broadcast prediction error, and offer testable predictions for future experiments.

## 1 Introduction

Almost three decades ago, Schultz et al. [1] proposed that midbrain dopamine (DA) neuron phasic activity encodes temporal difference errors (TDEs). This fundamental idea, leveraging the temporal difference (TD) learning algorithm [2], suggested the brain could assign credit in terms of expected future reward using temporally successive predictions. Numerous experiments have since substantiated this relationship within the TD reinforcement learning (TDRL) framework [3, 4, 5, 6, 7]. A core motivation was that engineered systems employ similar algorithms to optimise actions in complex environments, mirroring challenges faced by animals [1]. However, three key challenges currently impede TDRL research from fully realising this core motivation.

The first challenge arises because typical experiments and computational models, utilising single-attribute rewards (e.g., juice) and monolithic value functions, inadequately capture the multi-objective nature of challenges animals face in real life. Animals must constantly satisfy distinct, often conflicting objectives under changing priorities, a kind of non-stationary reward landscape. This necessitates either multiple value functions [8, 9] or alternative efficient representations like the successor representation [10, 11, 12], as standard RL approaches struggle with instant revaluation under non-stationary rewards [8, 9, 13]. This need for fast adaptation to multiple objectives (often encoded by different rewarding attributes), under shifting priorities, is ubiquitous in homeostasis [14, 15, 16] and extends to human cognitive tasks [17]. Neural evidence further suggests dopaminergic circuits projecting to different targets may encode multiple value functions corresponding to various reward modalities (e.g., food, juice, water [18, 19, 20, 21, 22]), valence [23], substance type [24], or even abstract features and contexts [25, 26, 27], yet this multiplicity is not captured by standard TDRL.

The second challenge concerns the TD-learning rule used by Schultz et al. [1], which learns values under the current behavioural policy (on-policy algorithms), rather than under an optimal policy (off-policy algorithms). The result is that the learned values estimate future returns assuming continued use of the current, often suboptimal, policy. While a core motivation was to learn optimal actions [1], on-policy algorithms characteristically learn suboptimal values under an often-exploring, suboptimal policy. This compromise, well-noted in traditional single-objective RL [28] and usually dealt with by heuristically tuning exploratory noise, becomes particularly problematic in multi-objective RL, where the behavioural policy can change drastically with shifts in needs or non-stationarity [8].

The third challenge stems from widespread dopamine heterogeneity, which calls into question the physiological basis of a single, broadcasted scalar reward prediction error (RPE). While a final scalar variable is computationally necessary to guide choice, the notion of a monolithic RPE signal is increasingly at odds with observations of diverse dopamine responses, both between and within striatal subregions. Extending the classical model [1] to multiple objectives [9] only partially explains this diversity (e.g., between different DA targets). For instance, within-target heterogeneity in regions like the tail of the striatum (TS) remains difficult to reconcile, as evidence points to a mix of threat and action prediction errors [23, 29, 30, 31, 32] that do not neatly integrate into the standard TDRL framework.

While distinct, the first two challenges—the need for multi-objective learning and the pitfalls of on-policy methods— converge on a single, fundamental question of optimal composition of multiple values. The brain clearly possesses mechanisms for learning multiple values, but how can they be reliably combined to produce a coherent and optimal policy? Ideally, this “recipe” for combining values should satisfy two key properties: optimality and compositionality. Optimality, the bedrock of modern reinforcement learning [33], simply means choosing the best possible action [34]. Compositionality, conversely, is the formal principle of constructing solutions to complex problems from a set of modular components, for instance by allowing a multi-attribute task to be represented by a basis set of value functions tuned to individual reward dimensions. Crucially, this structure allows policies for novel priority landscapes to be constructed by flexibly recombining these components, obviating the need to learn each new configuration de novo.

As noted previously [7], the choice of learning algorithm is not a mere technicality but a foundational choice dictating the system’s computational objectives. Recent proposals often overlook this, foregoing either optimality [9] or composability [8]. On-policy methods (e.g., SARSA), for instance, suffer from learning interference, especially under shifting contextual priorities. When the current context prioritises one reward, the agent’s policy becomes biased; because learning is tied to this policy, the valuation of alternative rewards is not learned in isolation but is corrupted by being evaluated through the lens of the current trajectory. In contrast, off-policy methods (e.g., Q-learning) fail because of the non-linearity of the Bellman optimality (max) operator, which is not additive. This creates an unrealistic assumption: the agent is treated as perfectly rational during valuation (identifying the single best future action) yet as boundedly rational during action selection (making stochastic choices). This inconsistency between ideal valuation and bounded action selection corrupts the composition of different reward values.

To resolve this, we adopt a normative framework from control engineering [35, 36, 34, 37], broadly termed linear RL [38]. We propose that the dopamine system’s objective is not merely to optimise cumulative reward, but to optimise returns augmented by a penalty for deviating from a default policy. This single modification provides a principled solution to the optimal composition problem [39]. By enforcing a consistent assumption of bounded rationality throughout both valuation and action selection, it resolves the paradoxical logic of standard algorithms, allowing multiple values to be robustly combined.

Remarkably, the same principle that ensures optimal composition provides a natural framework for understanding dopamine heterogeneity. The framework’s parallel architecture for outcome-specific prediction errors explains between-target dopamine heterogeneity [9, 40] and supports efficient learning across multiple rewards. We substantiate these claims in our Results, which use a didactic example and simulations to demonstrate these normative advantages across multi-objective tasks. Lastly, we discuss how this compositional logic can be extended to explain within-target heterogeneity, formally reconciling threat and action prediction errors in the TS.

## 2 Results

### 2.1 Theory sketch

At Marr’s computational level [41], we formalise dopamine’s objective as maximising future returns augmented by a penalty for deviating from a default policy. The agent thus optimises a relative-entropy–regularised return,

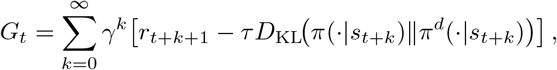

where *D*_KL_ is the KL divergence between the current behavioural policy *π* and a default policy *π*^*d*^. If *π*^*d*^ is uniform, this encourages random exploration (maximum-entropy RL); if *π*^*d*^ slowly tracks the learned policy *π*, it promotes choice perseveration or soft-habit formation by penalising deviations from recently taken actions [42, 38, 43, 44]. Lastly, we note that entropy-regularised RL is mathematically equivalent to planning/control as probabilistic inference [45].

At the algorithmic level, this objective is solved using soft Q-learning [46], an off-policy temporal difference (TD) method, adapted here for the general relative-entropy objective. Q-values are updated as per,

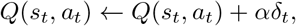

where *α* is the learning rate, and *δ*_*t*_ is the temporal-difference error at timestep *t*, defined as:

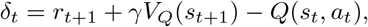

where the soft state-value is given by *V*_*Q*_(*s*) = *τ* log *E*_*a*_ *∼* _*π*_*d* exp(*Q*_*π*_(*s, a*)*/τ*) (see Methods, Eq. 7). *V*_*Q*_(*s*) replaces the biologically intractable max_*a*_ *Q*(*s, a*) of the standard Bellman optimality operator with a smoother, compositional form. Importantly, since the valuation of the next state is dependent on the default policy, separate from the current behavioural policy, soft Q-learning is an off-policy algorithms.

Notably, for this one-step algorithm, the KL-divergence from the objective does not appear in *δ*_*t*_ because the action *a*_*t*_ has already been taken [47, 46, 48], however it appears in eligibility trace-based extensions [49, 48]. Actions are chosen using a Boltzmann policy that is optimal under this regularised objective (Methods, Eq. 6).

Multiple soft Q-functions *Q*_*i*_(*s, a*) are learned in parallel for distinct reward attributes *r*_*i*_ and then combined through a weighted softmax or summation operation

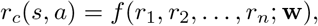

where the weights **w** reflect current priorities, such as homeostatic needs [14, 9], inferred beliefs [25, 26], or survival imperatives. When *f* is weighted softmax, the compositionality of optimal control laws in linear MDPs [39] guarantees that composing these independently learned values using the same function *f* yields a policy that is optimal for the composite reward *r*_*c*_.

At Marr’s implementation level, our model predicts parallel soft Q-learning mechanisms to account for heterogeneity in dopaminergic responses between distinct neural targets (Fig. 1). Separate value channels for distinct reward attributes support efficient multi-attribute learning while preventing positive outcomes from overriding threat or punishment values [50, 51, 52].

**Figure 1.**
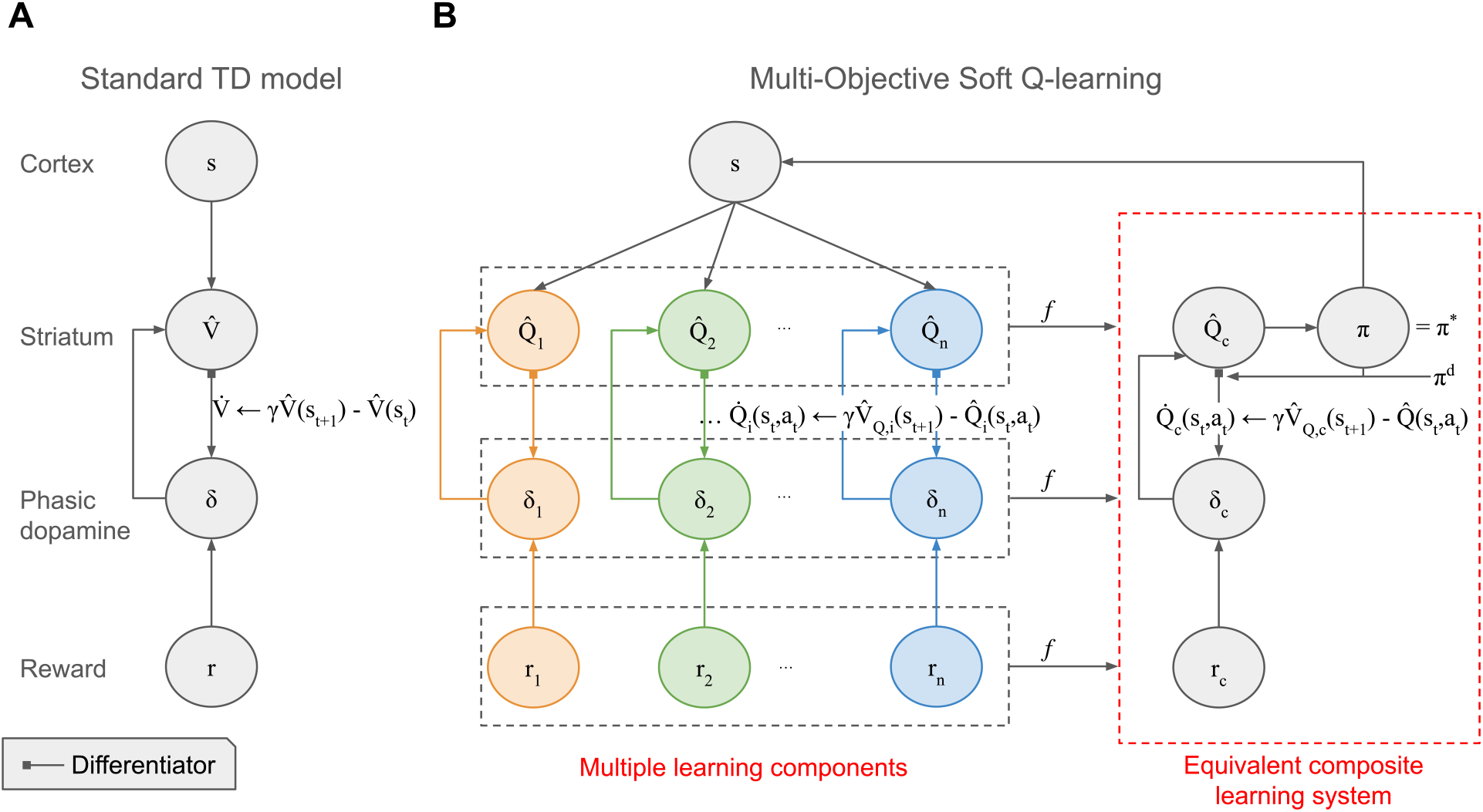
Neural architecture for optimally composable multi-objective reinforcement learning. (A) Conventional temporal-difference (TD) model [1]: dopamine neurons compute a scalar reward-prediction error (RPE) that updates a single state-value function in the striatum. Extensions with parallel outcome channels yield vectors of values and RPEs [9]. (B) Proposed model: parallel soft Q-learning modules learn distinct state–action values within a linearly solvable MDP framework [36]. Their outputs are optimally composed (*f*) to guide behaviour that maximises a weighted combination of rewards. This architecture supports parallel dopaminergic targets, requiring access to a default policy *π*^*d*^ for computing soft state-values *V*_*Q*_.

We first illustrate the principles of optimal value composition with a didactic simulation, building on Todorov [39], while situating and comparing with contemporary biological multi-objective RL models, then proceed to more concrete experiments.

### 2.2 Optimal composition of multiple value functions for reliable optimisation

A central challenge in multi-objective reinforcement learning (RL) is to reliably combine multiple value functions into a single, coherent policy. Recent proposals typically linearly decompose a composite reward *r*_*c*_ into attributes (*r*_1_, *r*_2_, …), learn independent value functions *Q*_*i*_(*s, a*) for separate reward components *r*_*i*_, and combine them linearly to form a composite value *Q*_comp_ = ∑ _*i*_*w*_*i*_*Q*_*i*_ [53, 8, 9]. However, depending on the specific value learning algorithm, this can lead to a critical trade-off: either the individual values *Q*_*i*_ are optimal for their respective rewards *r*_*i*_ but their composition *Q*_*comp*_ is not optimal for *r*_*c*_, or the composition is well-defined but the individual values *Q*_*i*_ themselves are sub-optimal.

To illustrate the first issue—individually optimal values that compose sub-optimally—we consider the off-policy, multi-objective Q-learning devised by Russell and Zimdars [53] and utilised by Dulberg et al. [8]. This method learns optimal *Q*_*i*_ for each *r*_*i*_ using standard Q-learning and then additively combines them. In a two-step MDP with two reward functions *r*_1_, *r*_2_ (Fig. 2A), if we equally weight rewards (*w*_1_ = *w*_2_ = 0.5) to get *r*_*c*_, the (undiscounted) optimal Q-values for *r*_*c*_ in the starting state *S*_0_ are 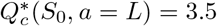 and 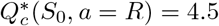 However, the additively composed Q-values are *Q*_*comp*_(*S*_0_, *a* = *L*) = *Q*_*comp*_(*S*_0_, *a* = *R*) = 5. Consequently, an agent using *Q*_*comp*_ with a softmax policy chooses actions L and R with equal probability, irrespective of the temperature *τ*, deviating from the optimal behaviour for *r*_*c*_ (Fig. 2B). This sub-optimality arises from the non-linearity of the max operator in the Bellman optimality equation.

**Figure 2.**
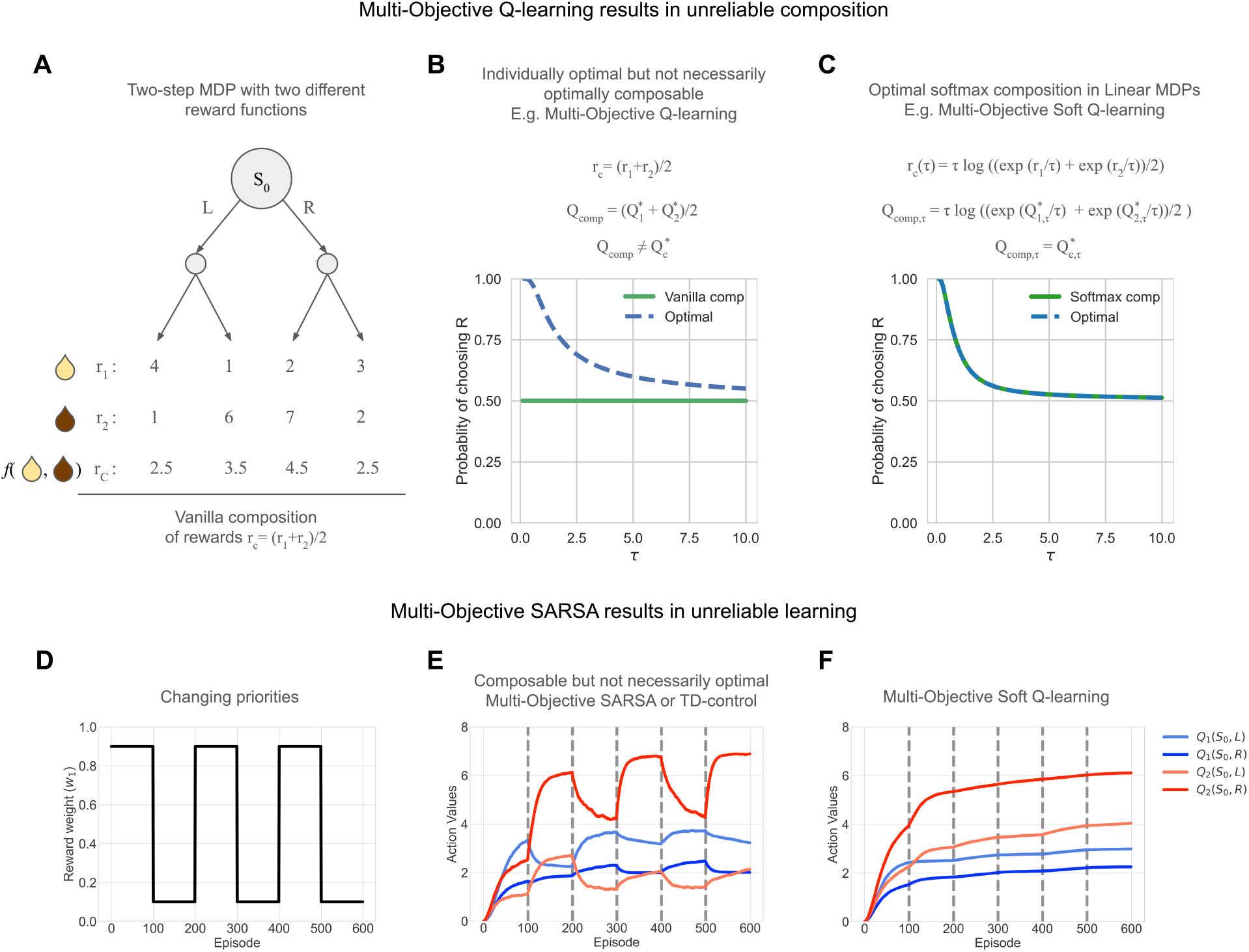
Demonstration of the reliable and optimal composition of values in linear MDP. (A) Two-step MDP with two (diverging) reward functions, say Juice 1 (*r*_1_) and Juice 2 (*r*_2_). (B) Action selection probabilities under sub-optimal additive composition of Q-values in MDPs (*Q*_*comp*_) deviate from optimal behaviour mandated by 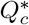 optimising *r*_*c*_. (C) Action selection probabilities under optimal softmax composition in Linear MDPs (*Q*_*comp*_) match optimal behaviour mandated by 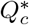 optimising *r*_*c*_. Weights set to *w*_1_ = *w*_2_ = 0.5 (equal priority for both rewards) and action probabilities plotted for a range of *τ*. (D) Changing reward weight *w*_1_, denoting the change in priorities and *w*_2_ = 1 − *w*_1_. (E) Action values of multi-objective (MO) SARSA show unstable and unreliable learning over episodes. (F) Action values of MO Soft Q-learning show reliable and stable learning over episodes. Note that the focus is on value learning over episodes is stable or has interference; action values from (E) and (F) cannot be compared as the objective functions are different.

In contrast, our approach using soft Q-learning within the linear MDP framework allows for a reliable composition. Here, rewards *r*_*c,τ*_, a composite reward function composed from individual rewards *r*_1_ and *r*_2_, and Q-values *Q*_*comp,τ*_ are functions of *τ*, which controls the influence of a default policy *π*^*d*^ (here, uniform). As shown in Fig. 2C, the resulting composed policy reliably optimises the target reward composition, achieving 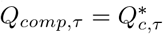 (Theorem 1, Methods Section 4.3). This demonstrates optimal value composition [39].

On the other hand, works that employ on-policy algorithms, i.e., TD(0) [1], its multi-objective extension [9], and extensions to control (SARSA), are known to not reliably learn optimal policies, especially from sub-optimal trajectories (see supplementary Fig. S2).

The limitations of on-policy learning become particularly acute in multi-objective scenarios with dynamically shifting priorities. This is because policies optimal to one value component are bound to be sub-optimal for other value components, but values for all components are learnt under a common behavioural policy. Here, we highlight a critical form of interference in multi-objective (MO) on-policy algorithms such as MO SARSA (or vanilla Reward Bases [9]): as priorities shift (Fig. 2D), the ensuing changes in the global behaviour policy directly impact the valuation of individual components (supplementary Fig. S1 illustrates different policies under different priorities). Because on-policy value updates depend on the current policy *π*, actions taken to optimise one reward modality can lead to unintended revaluation and even unlearning of values for other modalities (Fig. 2E). The unstable value estimates demonstrate how an on-policy agent struggles to adapt: its learned value for ‘Juice 1’ is also inadvertently altered while it pursues ‘Juice 2’, preventing a truly flexible switch when the rules change. In contrast, our off-policy multi-objective (MO) soft Q-learning framework effectively mitigates this interference, as the valuation of the next state is dependent on the default policy separate from the current behavioural policy. This ensures stable learning of all value components (Fig. 2F).

### 2.3 Efficient learning and off-policy fast adaptation

Efficient adaptation to non-stationary rewards is a hallmark of intelligence. While full model-based RL offers a solution, it is computationally costly. Part model-free solutions rely on either (i) composing independent value functions for different (pre-defined) reward types, as discussed above [8, 9], or (ii) learning efficient representations such as the successor representation (SR) [10], which allow rapid revaluation of policies when reward functions change.

Crucially, in the case of on-policy model-free algorithms, Millidge et al. [9] show that a combination of TD(0) learning rules for each of the reward types is akin to a compressed SR with rewards tuned only to relevant dimensions. An equivalent relationship in off-policy algorithms is lacking. The default representation (DR), an off-policy counterpart to the SR derived from linear MDPs [38, 36, 34], overcomes the on-policy limitations of the SR.

#### Theoretical result: Relationship to the default representation (DR)

MO Soft Q-learning learns values equivalent to those learned by a compressed DR tuned to only relevant (predefined) reward dimensions (Methods, Section 4.5). It further provides two benefits: First, MO Soft Q-learning scales linearly with state space size (assuming a fixed, smaller number of reward dimensions), whereas the full DR scales quadratically. Second, unlike the SR, the DR lacks an efficient TD learning algorithm and typically requires matrix inversions for its computation. MO Soft Q-learning provides a TD-based mechanism to learn DR-like values.

#### Simulation result: MO Soft Q-learning enables superior adaptation to shifting priorities

We empirically tested these advantages in a four-room grid world where an agent pursued one of three goals, with priorities shifting every 1000 episodes (Fig. 3A; see Methods for more details).

**Figure 3.**
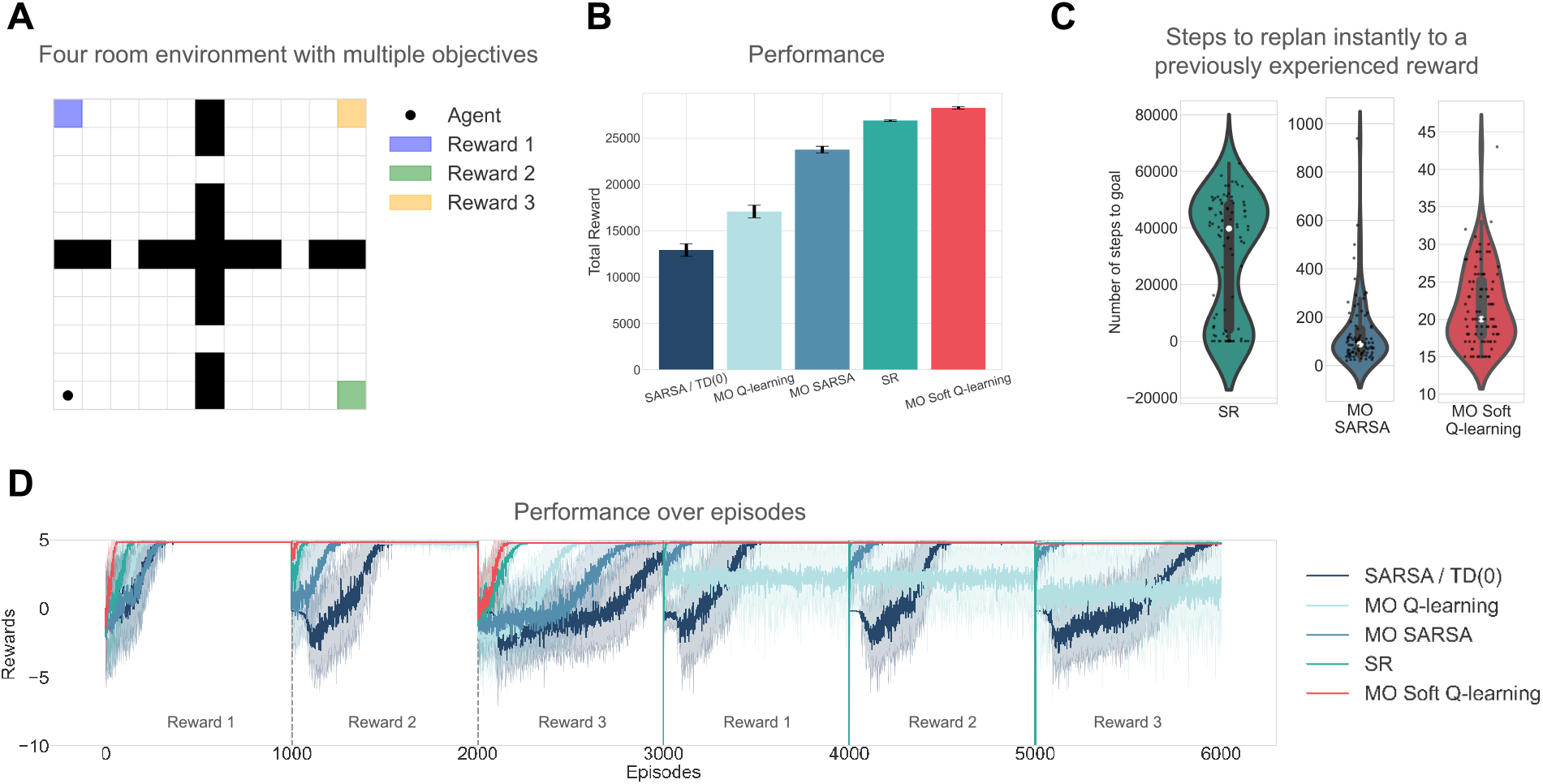
Demonstration of efficient learning and fast adaption to changing priorities in a four-room environment. (A) Four-room environment with changing priorities between three rewards every 1000 episodes, and the agent starts in the same starting position. Rewards for goals: +5 points, step cost: -0.01. The episode terminates only on reaching one of the rewarding goals. Meta-parameter *τ* = 0.5 to allow all algorithms to converge to an efficient path. (B) Multi-objective (MO) Soft Q-learning algorithm performs the best amongst comparisons in terms of total rewards accrued, highlighting its fast adaptation capabilities. (C) Steps to the goal on episodes 3000, 4000 and 5000 are plotted for different algorithms to test replanning to a previously experienced reward. SR performs the worst at replanning, requiring substantial policy re-evaluation, while MO Soft Q-learning performs the best. (D) Performance over episodes shows different rates of adaptation for different algorithms upon a change of priorities.

MO Soft Q-learning outperformed SR and other MO TD algorithms in total rewards accrued (Fig. 3B), demonstrating superior adaptation. The policy-dependence of SR was particularly detrimental during priority shifts requiring substantial re-planning [54]. For instance, when needing to switch back to a previously learned goal after extensive training on another, the SR agent took orders of magnitude more steps than MO Soft Q-learning or even MO SARSA (Fig. 3C). MO Soft Q-learning, by maintaining relatively stable, independent values for each reward, could immediately leverage the appropriate value function. The adaptation rates over episodes further illustrate these differences (Fig. 3D).

Lastly, we observe that off-policy algorithms continue to propagate optimal Q-values for all components, while collecting data under different policies (priorities), unlike on-policy algorithms (supplementary Fig. S4). However, MO Q-learning requires commensurate temperature annealing to get the most benefits, whereas MO soft Q-learning manages this trade-off without explicit annealing.

In summary, dopaminergic circuits may cache outcome-specific value functions that can be flexibly combined according to changing physiological or contextual priorities. These functions can be mapped to different DA targets, responsible for different reward bases (for example, see Millidge et al. [9]).

## 3 Discussion

In this work, we address the fundamental challenge of reliably optimising multiple objectives in reinforcement learning and propose a model that outperforms existing multi-objective methods. This invites a reconsideration of the dopaminergic system’s computational objective, shifting from the classical view of cumulative discounted reward maximisation [1] towards optimising returns augmented by a KL penalty for deviations of the behavioural policy from a default policy. This reframing not only yields distinct functional advantages but also generates novel testable predictions in efficient and stable learning. In doing so, we also then discuss how the same framework can be instantiated to address conflicting views of TS-projecting dopamine in threat prediction errors (TPEs) and action prediction errors (APEs) within the temporal difference reinforcement learning framework.

This approach was designed to directly address several limitations of the standard model, specifically the challenge of flexibly pursuing multiple, often conflicting, rewards without the learning process becoming unstable or inefficient—a known issue for classic on-policy methods. Our model’s core innovation is to augment the classic reward prediction error with a regularisation term that imposes a ‘cost’ for deviating from a default policy.

At an algorithmic level, this work addresses how to ensure (often individually optimal) value functions cooperate effectively to drive reliable behaviour, despite competing for control. This contrasts with “delegation” approaches [55, 56, 57], where only one value function controls actions at any timestep, thus avoiding this problem. Regarding our optimal composition results, previous multi-objective TD(0) or SARSA approaches that scale on-policy prediction errors with weights (e.g., state-dependent RPE modulation [9, 58] or feature-specific weight updates [26, 40]) may slightly ameliorate unreliable learning or policy interference, but do not resolve it (supplementary Fig. S3). This excludes trivial conditions where only one value component is active, preventing interference. A better alternative for these models might be to use importance sampling, which would ensure off-policy learning.

Off-policy learning is a prominent theme in this work, as it demonstrably improves performance over on-policy multi-objective RL algorithms and the successor representation, while also relating directly to its off-policy counterpart [38]. The brain may implement off-policy learning for several reasons: First, it prevents interference and unlearning between multiple values amidst changing priorities. Second, it facilitates learning amidst motor noise and competition from distributed control systems, like the motor cortex and cerebellum [59, 60]. Third, on-policy algorithms, such as the successor representation, exhibit strong policy dependence where goal information contaminates the state map, hindering flexible transfer [54, 61], a problem solved by off-policy alternatives [38]. Fourth, the ability of episodic memories to utilise cached values [62, 63] or stale behavioural data for performance improvements points towards an underlying off-policy mechanism, akin to its necessity in deep Q-learning’s episodic replay buffers [64]. However, finding strong neural and behavioural evidence for interference and unintended unlearning between two or more value systems (for different rewards) under changing priorities in a two-step task similar to Fig. 2D-F, would falsify our hypothesis of phasic dopamine performing off-policy multi-objective RL and find evidence for on-policy multi-objective RL (e.g. [9]). Rapid change in priorities could be potentially implemented as a task rule that needs to be inferred.

Our framework synthesises several threads of the evolving dopamine story, which has progressively expanded from a simple scalar reward signal [1] to a multifaceted control signal. Our results (Fig. 3C) also highlight the behavioural inefficiencies of the SR model in replanning after overtraining [54]. Indeed, SR models of dopamine often need to treat reward as a feature to explain RPEs as a form of state prediction error [12], as Lee et al. [40] shows they otherwise fail to consistently respond to rewards. This highlights the benefit of outcome-specific models such as ours [and that of 9], which efficiently achieve values comparable to an SR/DR model [10, 38] when tuned to a subset of reward types.

In terms of limitations; first, though broadly applicable to decision-making under changing priorities, when applied to homeostatic priorities, it cannot explain physiological state-dependent modulation of prediction errors [65]. This is a limitation common to all multi-objective RL approaches (e.g., those by Dulberg et al. [8], vanilla version of Millidge et al. [9] and ours), which learn equally from all reward types, at all times. Second, despite its advantages, soft maximum composition optimises an objective different from a simple weighted sum of utilities. While this is not necessarily an issue for modelling homeostasis, where value representation is debated [15, 66, 8], and may even account for inhibitory effects of irrelevant drives [15, 67], we find it can fit poorly to human behaviour when participants explicitly maximise weighted sums of utilities or show generalisation in task structure rather than in values [17] (see supplementary Fig. S5). Third, while there are compelling reasons for the brain to implement off-policy learning [59, 60, 32], some early work [68, 69] suggested phasic dopamine implements on-policy algorithms like SARSA. However, those experiments involved overtrained monkeys where no learning occurred, making observed TD error signals potentially epiphenomenal. Our framework could partly explain these differences in action values as the KL divergence from the overtrained default policy (APEs). Furthermore, the observed use of cached values in computing TDEs [62, 63] better aligns with off-policy rather than on-policy TD learning.

Finally, while this paper focuses on the reliable composition of distinct reward attributes, the same linear MDP framework may also accommodate within-target dopaminergic heterogeneity. As a discussion-level illustration of this broader scope, Fig. 4 summarises a TS-focused instantiation in which threat prediction errors and action prediction errors can coexist within a single entropy-regularised TD-error framework [29, 30, 32]. The detailed model, simulations, and experimental predictions are developed in related work [70]; here, the example is included only to show how the same computational objective can extend beyond between-target value composition. The TS-focused variant extends soft Q-learning with eligibility traces, resulting in: (1) a more biologically plausible computation of the TD error using the temporal derivative of the soft Bellman value function; (2) the introduction of a KL term in the TD error, which shows features similar to action prediction errors (APEs); and (3) the use of APEs [32, 42, 38] to update the default policy. We explore these TS-specific implications and their normative benefits for cautious exploration and stable learning under uncertainty in related work.

**Figure 4.**
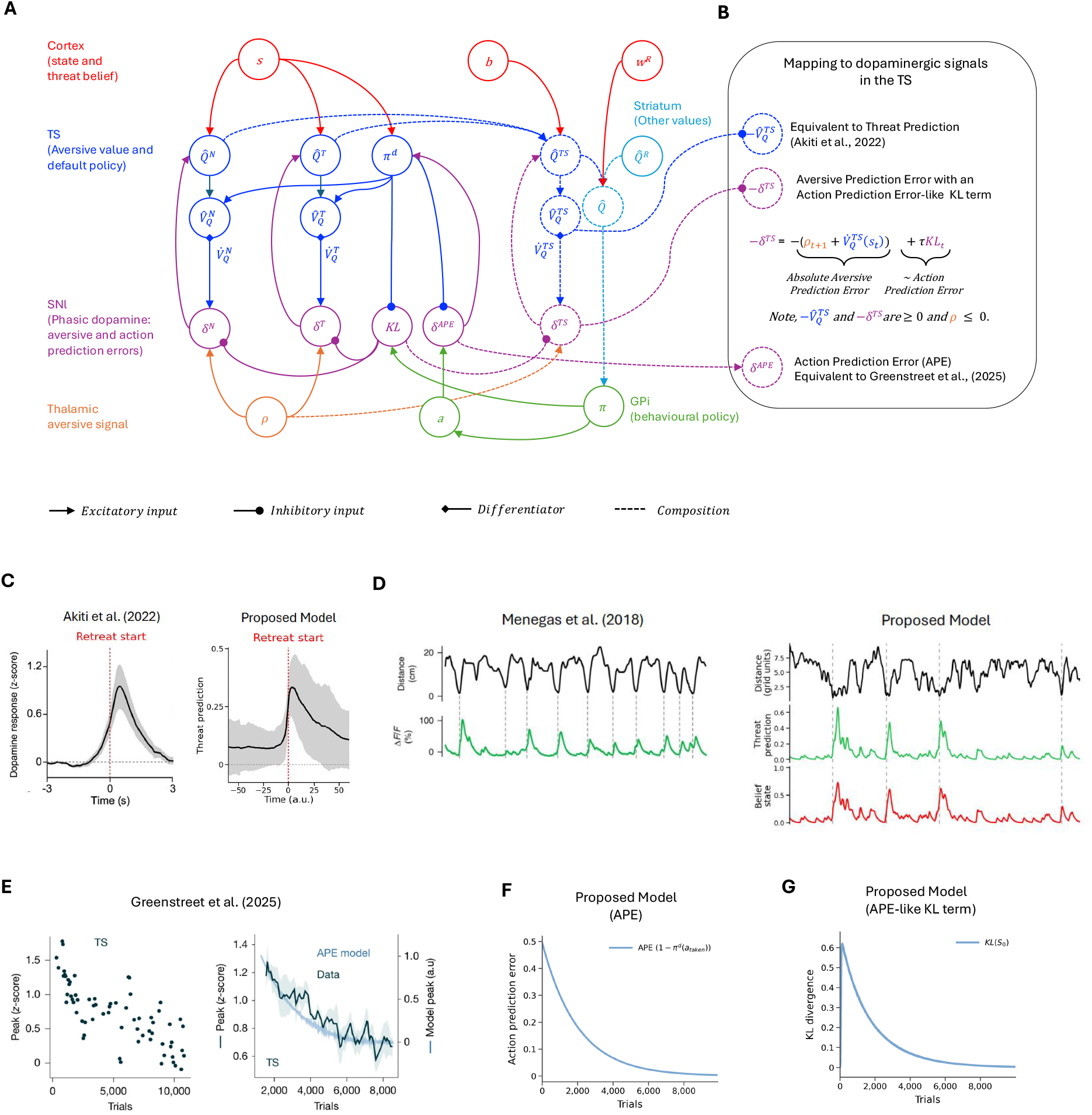
Discussion-level summary of a TS-focused instantiation of the framework [70]. (A) A colour-coded wiring diagram of the proposed TS model. 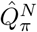 and 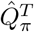 are Q-value functions for the ‘Not-threatened’ and ‘Threatened’ contexts, respectively, while 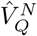 and 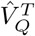 are the corresponding soft Bellman value functions. 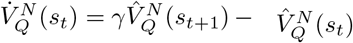 and 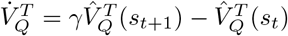 are temporal derivatives of the soft Bellman value function. *ρ* is the aversive feedback signal *ρ* = min(*r*, 0) where *r* ∈ (−∞, + ∞), shared between the value functions in both contexts, contributing to the aversive TD errors 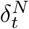 and 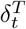. The default policy is updated based on the chosen action using action prediction errors [32, 42], scaled by the learning rate *α*^*d*^, and then normalised. The aversive values are updated by scaling the aversive TD errors by the learning rate *α* and eligibility traces *e*_*t*_(*s, a*) [49, 48]. The TS Q-values are composed using weights derived from a belief state *b*, and can be further integrated with other reward-based value functions according to their respective weights *w*^*R*^. (B) The mapping of model signals to the dopaminergic signals observed in experimental evidence, and to previous models that attempt to explain them [30, 32]. TD-error equation suggests why an aversive prediction error and an action prediction error-like KL term may co-exist in the TD errors, alongside previously proposed threat prediction (TP) values and APEs. (C) In the TS-focused model instantiation, Threat Prediction qualitatively reproduces the TS activity observed after the start of a retreat but not during approach [30]. (D) The same result is plotted along a continuous timeline before averaging, alongside the data from Menegas et al. [29]. Δ*F/F* represents the TS activity. The proposed model demonstrates higher TP signals at the start of a retreat (denoted by a dotted line). (E) Greenstreet et al. [32] observe that TS activity decreases across trials as mice acquire soft habits, which they qualitatively model using action prediction errors (APEs) that drive “value-free” habits [42]. (F) In the TS-focused model instantiation, the APEs used to update the default policy in our model (also in a value-free manner) replicate the APE curves from Greenstreet et al. [32]. (G) The KL term in our proposed model initially shows an increase and then an APE-like decrease after a few early trials. All figures are retrieved and minimally adapted from Menegas et al. [29] with permission from Springer Nature and from Akiti et al. [30] under [CC-BY-NC-ND 4.0](https://creativecommons.org/licenses/by-nc-nd/4.0/deed.en) license with permission from Elsevier. Figures adapted from Greenstreet et al. [32] under a [CC-BY 4.0](https://creativecommons.org/licenses/by/4.0/deed.en) license (figures were cropped and combined).

In offering a novel normative framework for multi-objective reinforcement learning, this paper re-conceptualises the computational role of striatal dopamine. Our findings demonstrate how the brain might achieve efficient and stable learning under changing priorities, while suggesting how the same framework may extend to cautious behaviour and within-target dopamine heterogeneity.

## 4 Methods

### 4.1 Reinforcement learning in MDPs

Let the environment be a Markov Decision Process, where at time *t* = 0, 1, 2, …, the agent is in state *s*_*t*_ ∈ *S* and takes action *a*_*t*_ ∈ *A* and receives the next state *s*_*t*+1_ ∈ .*S* and the reward *r*_*t*+1_ = *r*(*s*_*t*_, *a*_*t*_) ∈ *R*. The dynamics of MDP are given by the conditional probability *p*(*s*^*′*^, *r*|*s, a*) = Pr(*s*_*t*_ = *s*^*′*^, *r*_*t*_ = *r*|*s*_*t−*1_ = *s, a*_*t−*1_ = *a*).

The discounted return at time *t* is given by 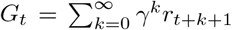 where *γ* ∈ [0, 1]. Policy *π*(*a*|*s*) is a mapping from states to the probabilities of choosing each possible action. The value function of a state *s* under the policy *π* is formalized as *V*_*π*_ = *E*_*π*_[*G*_*t*_|*s*_*t*_ = *s*], ∀*s* ∈ *S*. Similarly, the action-value function is *Q*_*π*_(*s, a*) = *E*_*π*_[*G*_*t*_|*s*_*t*_ = *s, a*_*t*_ = *a*].

The Bellman equation of a value function *V*_*π*_ is:

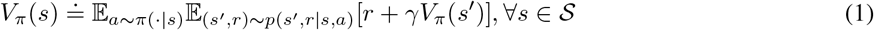

The Bellman optimality equation is:

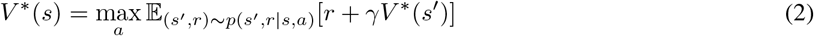

### 4.2 Entropy-regularised reinforcement learning in Linear MDPs

Entropy-regularised RL [36, 34, 71] augments the reward function with a term that penalises deviating from some default policy *π*^*d*^, essentially making “soft” assumptions about the future policy (in the form of a stochastic action distribution). When *π*^*d*^ is an uniform policy, this reduces to max entropy reinforcement learning [47, 46]. The expected reward on taking action *a*_*t*_ in state *s*_*t*_ is given by 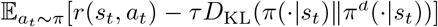, which can be further compactly written as 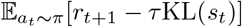. Here, *τ* is the scalar temperature parameter, and KL(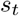) is the Kullback-Leibler divergence between the current policy *π* and a default policy *π*^*d*^ in state *s*_*t*_. Thus, the entropy-augmented return is 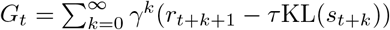

The value function definitions under a policy *π* at any timestep *t* based on the entropy-augmented returns are as follows,

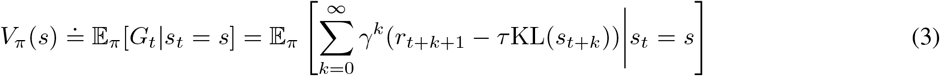

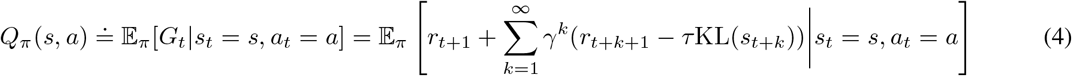

Note that this Q-function does not include the first KL penalty term (KL(*s*_*t*_)), as it does not depend on action action *a*_*t*_ which has already been chosen [47, 46, 48]. This gives the following relationship which holds for all policies *π*.

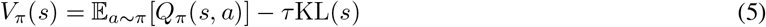

The greedy (stochastic) policy in entropy-regularised RL is the Boltzmann policy (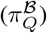):

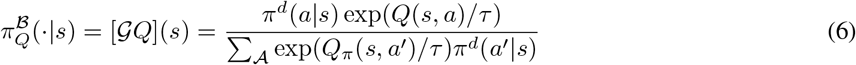

Under the Boltzmann policy, the Bellman equation is equivalent to the “soft” Bellman equation, performing a soft maximum operation over Q-values:

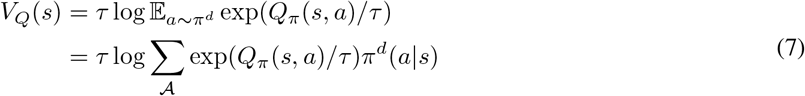

Note, this log-sum-exp performs a soft maximum because, max{*x*_1_, …, *x*_*n*_} ≤ softmax(*x*_1_, …, *x*_*n*_) ≤ max{*x*_1_, …, *x*_*n*_} + log(*n*).

### 4.3 Multi-objective reinforcement learning and optimal composition in Linear MDPs

Having discussed reinforcement learning (RL) in MDPs and Linear MDPs with single-attribute rewards, we now focus on multi-objective RL, which concerns multiple rewarding attributes **r** = [*r*_1_, *r*_2_, …, *r*_*n*_]. Note, taking an action at timestep *t* results in rewards *r*_*i,t*+1_, for the *i*-th reward dimension, but we will omit the time subscript from here for convenience. The objective is to maximise a cumulative discounted return of a reward function composed of these attributes:

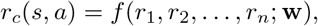

where **w** is a set of non-negative parameters that weight each attribute, satisfying ∑ *w*_*i*_ = 1. We address the problem of *optimal compositions*: determining how the reward function should be composed of multiple attributes to motivate meaningful behaviour and how to compose value functions to ensure the resulting policy acts optimally with respect to the composed reward function.

As shown in Results section 2.2, a simple linear composition of Q-values, such as 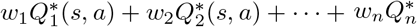 (*s, a*), may not maximise the composed reward function

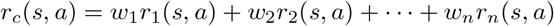

due to the non-linearity introduced by the *max* operation in the Bellman optimality equation in MDPs:

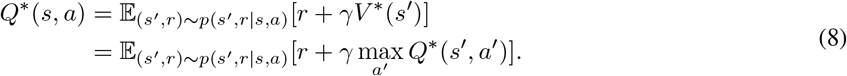

However, in linear MDPs [35, 36, 39, 34], which replace the *max* operation over Q-values by a soft-maximum *V*_*Q*_ with respect to the default policy (see equation 7), optimal (softmax) and near-optimal (additive) compositions are possible. The Bellman optimality equation for Q-values in linear MDPs is as follows:

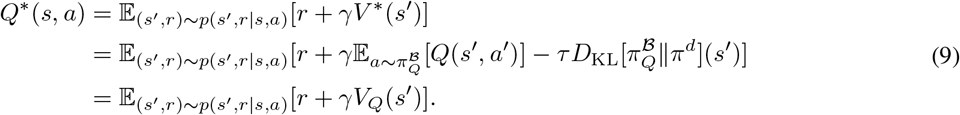

#### Theorem 1 (Optimal Softmax Composition) *[39, 71]*

*Let* 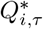 (*s, a*) *be the optimal entropy-regularized Q-functions for individual rewards r*_*i*_(*s, a*). *Then the reward function for the composed task is given by:*

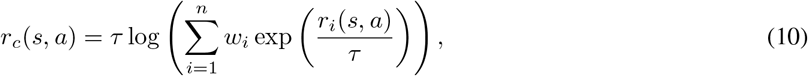

*The optimal Q-function* 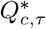 (*s, a*) *for the composed task is equal to the composition of Q-values Q*_*comp,τ*_ (*s, a*) *given by:*

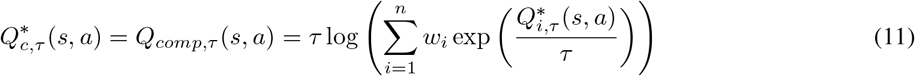

The theorem for additive composition in Linear MDPs is provided in the Supplementary Methods 2.

### 4.4 Off-policy model-free learning algorithms in Linear MDPs Soft Q-learning (One-Step)

We adopt soft Q-learning and extend it from the maximum entropy formulation to a relative entropy formulation. The Q-value update equation is:

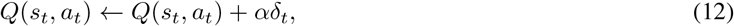

where *α* is the learning rate, and *δ*_*t*_ is the reward prediction error at timestep *t*, defined as:

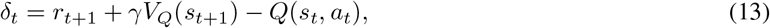

where *V*_*Q*_ is given by equation 7. For multi-step extensions involving eligibility traces please refer to Mahajan and Seymour [49], Schulman et al. [48].

### 4.5 Relationship to the default representation (DR)

The optimal values *V* ^*∗*^(*s*) are calculated using the default representation as follows [38, 36, 34] (in a model-based fashion):

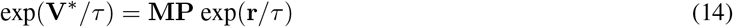

where, **V**^*∗*^ is the vector of optimal values at nonterminal states, **r** is the vector of rewards at terminal states, **P** is the one-step transition probabilities **T**_*NT*_ from non-terminal states to terminal states under the default policy *π*^*d*^ and **M** is the DR matrix defined as **M** = (diag(exp(−**r**_*N*_)*/τ*) − **T**_*NN*_)^*−*1^, where **r**_*N*_ is vector of rewards at non-terminal states and **T**_*NN*_ is the one-step transition probabilities between non-terminal states under the default policy *π*^*d*^. Further, [38] extend and define a more general version of the DR matrix *D* over all states (not just terminal states), as **D** = (diag(exp(− **r**_*A*_)*/τ*) − **T**)^*−*1^, where **r**_*A*_ is vector of rewards over all states and **T** are transition probabilities over all states under the default policy *π*^*d*^. Here, **M** is a sub-block of **D** and **T**_*NT*_ and **T**_*NN*_ are sub-blocks of matrix **T**. We observe that in both cases, it requires storing and/or learning the one-step transitions **T** under the default policy over all states, resulting in a *S* × *S* memory cost, where *S* is the size of the state space. Therefore, the memory cost of the DR scales quadratically with the size of the state space.

We propose our model (composition of multiple soft Q-learning rules) is intuitively akin to a compressed DR tuned to only relevant reward dimensions and converges to the same. To show this, we decompose the reward vector at terminal states, by assuming it to be composed using the optimal softmax composition [39], 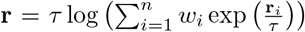. Therefore, the vector of optimal values using the DR will be:

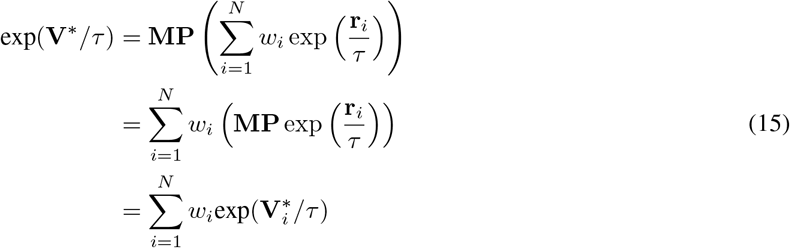

Here, the vector of optimal values for each reward decomposition can be computed using soft Q-learning and consumes memory cost of *S* (state-space size). Therefore, for *N* reward decompositions, our method consumes a memory cost of *S* × *N*. We believe the necessary reward decompositions would often be much lesser than the total number of states, *N << S*. Therefore, our method scales linearly with the size of the state space, whilst achieving the same optimal values as the DR for a predefined reward basis.

### 4.6 Simulation parameters

Simulations in Fig. 2D,E and supplementary Fig. S2, S3 use learning rate *α* = 0.1, discount rate *γ* = 1, temperature *τ* = 1 (unless explicitly varied) and are averaged over 100 runs. Simulations on efficient learning (Fig. 3) use *α* = 0.1, *γ* = 0.99, *τ* = 0.5 for all algorithms, and results are averaged over 30 runs.

## Acknowledgments

An earlier extended abstract of this work was presented at RLDM 2025, and the authors thank the two anonymous reviewers at RLDM 2025, Thomas Akam and Chris Summerfield for their helpful suggestions on how to make this work more impactful. Authors thank the funders: Wellcome Trust (214251/Z/18/Z, 203139/Z/16/Z and 203139/A/16/Z), IITP (MSIT 2019-0-01371) and JSPS (22H04998). This research was also partly supported by the NIHR Oxford Health Biomedical Research Centre (NIHR203316). The views expressed are those of the author(s) and not necessarily those of the NIHR or the Department of Health and Social Care. For the purpose of open access, the authors have applied a CC BY public copyright licence to any Author Accepted Manuscript version arising from this submission.

## Code and Data availability

Code for all simulations is available at https://github.com/PranavMahajan25/OptCompMultValues.git. No data was collected during this study.

## Supplementary Figures

**Figure S1.**
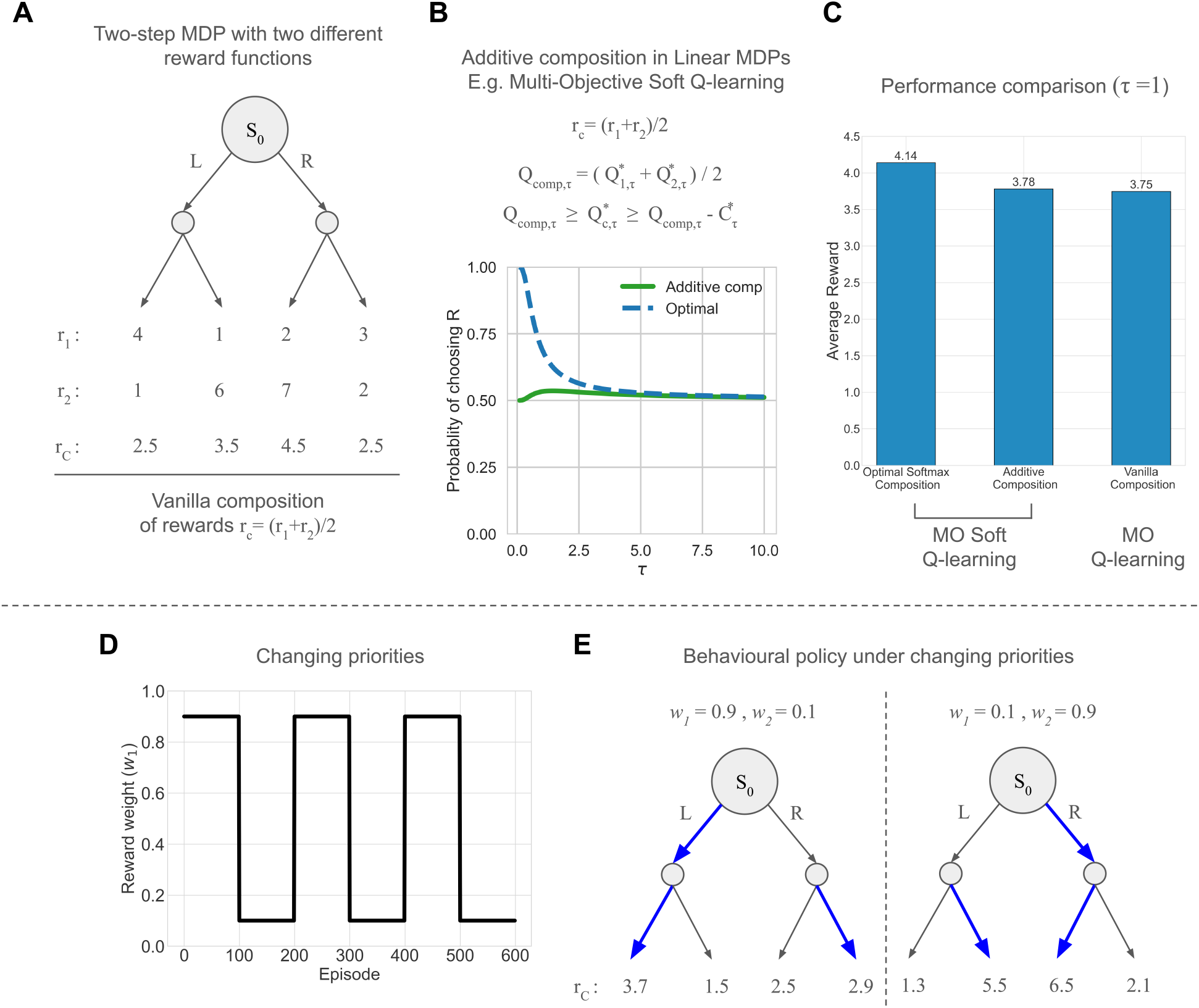
MO Q-learning comparison: (A) Two-step MDP with two (diverging) reward functions. (B) Action selection probabilities under additive composition in Linear MDPs. (C) Multi-objective (MO) soft Q-learning leads to better performance than MO Q-learning in this task. MO SARSA comparison: (D) Reward weight *w*_1_, denoting change in priorities. (E) Distinct behavioural policies under different priorities.

**Figure S2.**
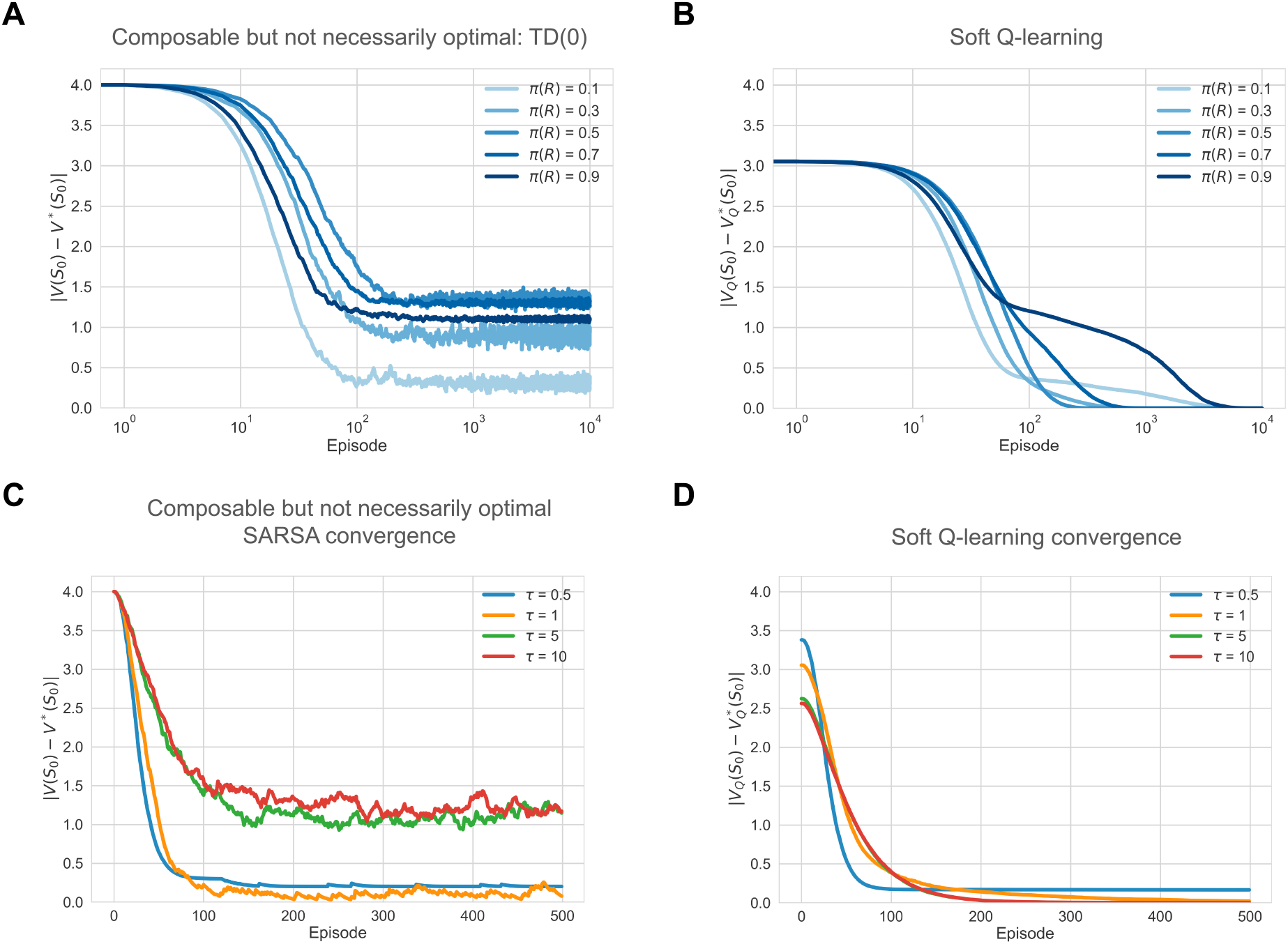
Demonstration of the off-policy learning of optimal values in linear MDP under sub-optimal trajectories (A & B) and under optimal control at different *τ* (C & D).

**Figure S3.**
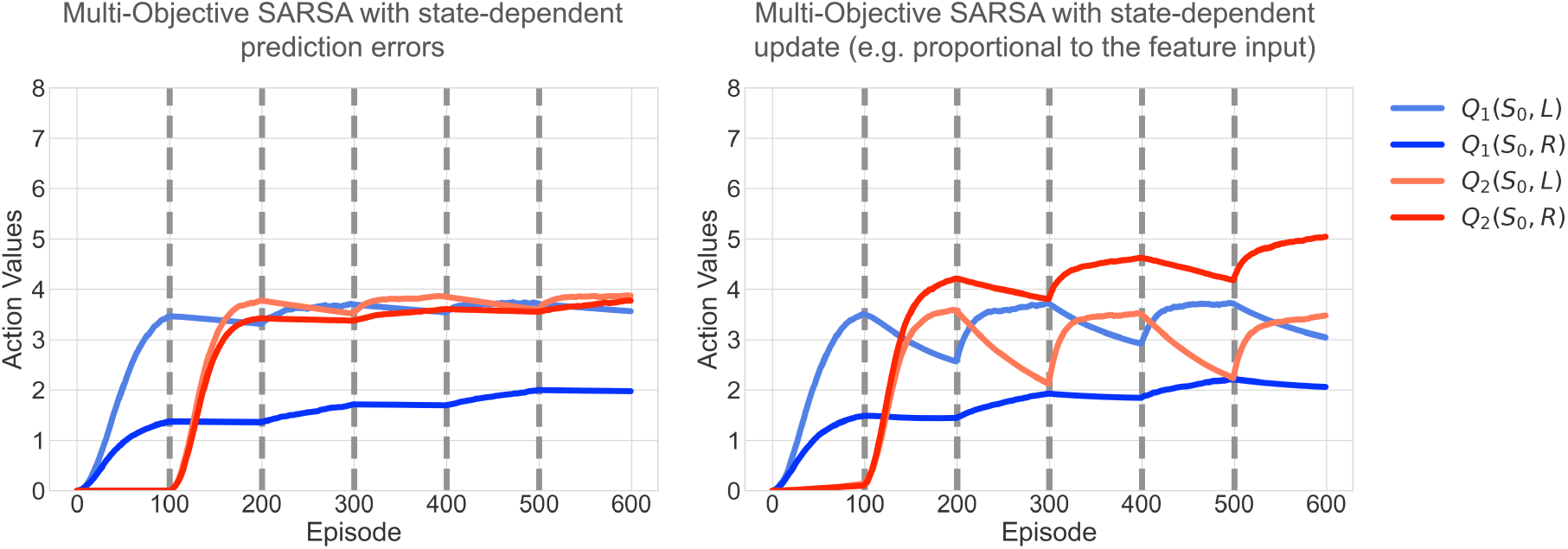
MO SARSA with state-dependent TD-errors, uses 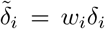. Neither variant completely avoids the interference and unintended unlearning effect caused by the on-policy nature of MO SARSA.

**Figure S4.**
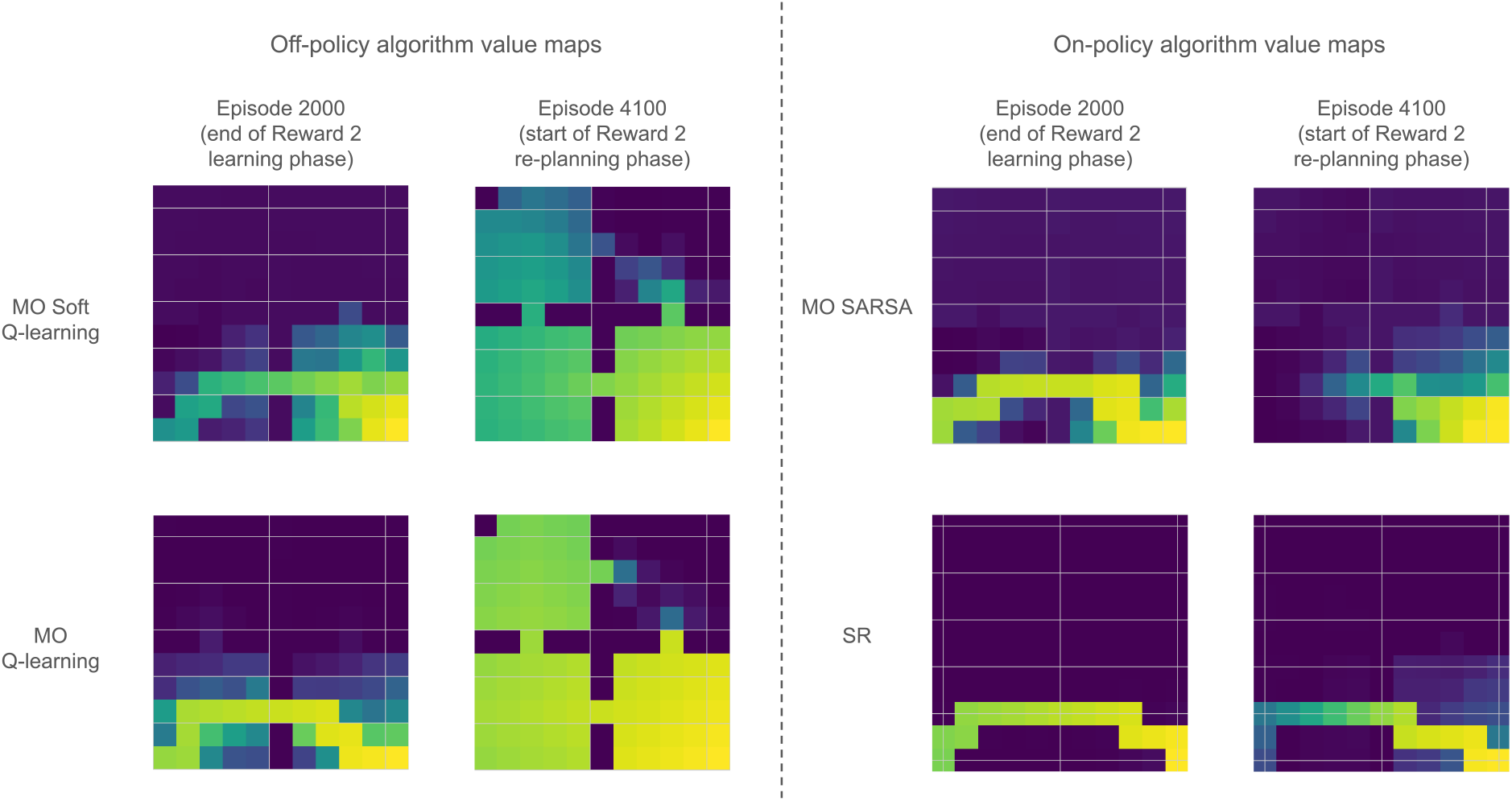
Differences in value propagation between multi-objective (MO) off-policy and on-policy algorithms, while exploring under different policies. Off-policy algorithms propagate optimal values throughout the environment, whereas on-policy algorithms do not.

**Figure S5.**
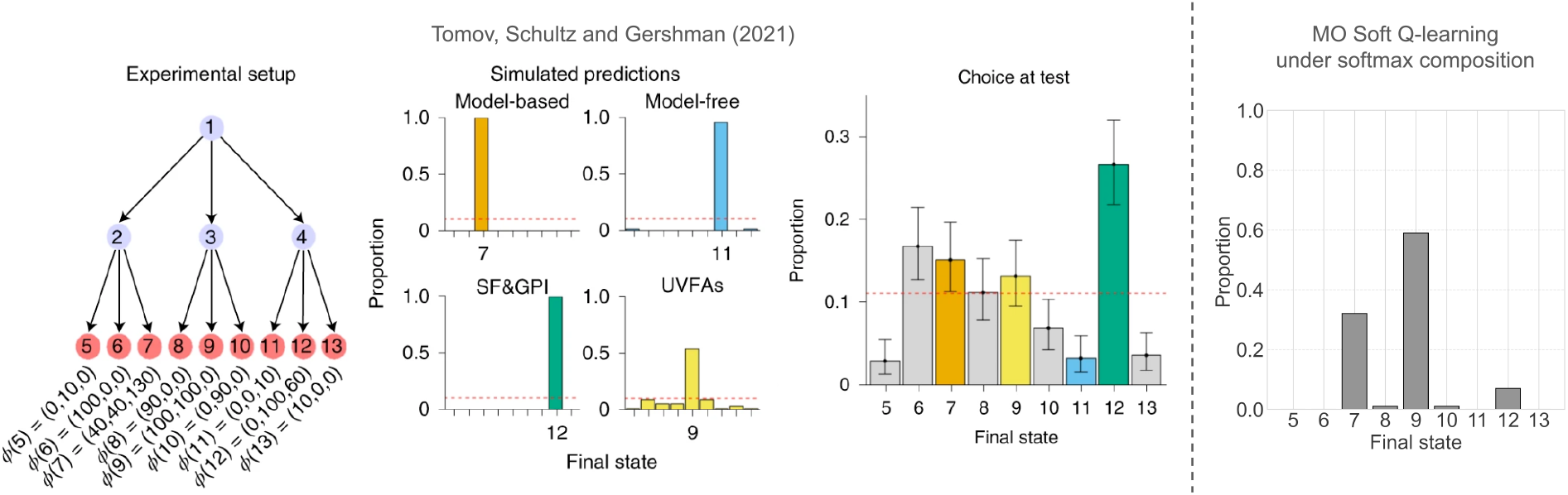
Soft maximum composition may not match human behaviour when they are explicitly asked to optimise the weighted sum of rewards across different modalities [17]. Figures adapted from Tomov et al. [17] with permission from Springer Nature.

## Supplementary Methods

### SM 1: How does the Boltzmann policy achieve the soft Bellman optimal policy in Linear MDPs?

We here aim to provide an intuitive explanation for known results. Consider the KL divergence between any policy *π* and the Boltzmann policy 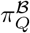 under some Q-values.

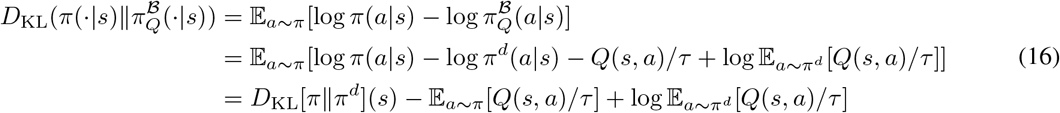

We can rearrange this equation and multiply by *τ* to get *V*_*π*_ (as per equation 5) on the left-hand side.

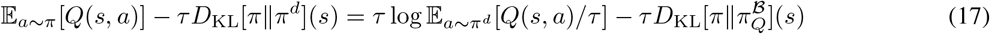

Here, we can see that the left-hand side of the equation (i.e. *V*_*π*_(*s*))) is maximised with respect to *π*, during generalised policy iteration (GPI), when the KL term on the right-hand side is minimized (as the other term does not depend on *π*), and 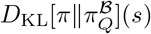 is minimized at 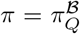. After each GPI, as 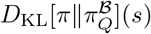 approaches zero at 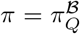, we observe that left-hand side of the equation is the “soft” Bellman value function *V*_*Q*_(*s*). This shows that for fixed Q-values, the Boltzmann policy is the stochastic greedy policy that maximises value. Under optimal Q-values, this greedy policy can lead to the Bellman optimal policy in linear MDPs.

During the generalised policy evaluation (GPE), these optimal Q-values can be learnt using any reinforcement learning algorithm with convergence guarantees. Repeating these generalised policy updates (GPE+GPI) will lead to the optimal policy *π*^*∗*^ will be given by the Boltzmann policy 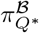. This concludes our intuitive explanation.

### SM 2: Theorem for additive composition in Linear MDPs

#### Theorem 2 (Additive Composition) *[72, 71]*

Let 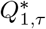 (*s, a*) *and* 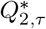 (*s, a*) *be the optimal entropy-regularized Q-functions for two tasks with rewards r*_1_(*s, a*) *and r*_2_(*s, a*).

*Then the reward function for the composed task aimed to ensure both objectives are given by the average of the individual reward functions:*

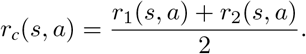

*Let the composition of Q-values Q*_*comp,τ*_ (*s, a*) *be:*

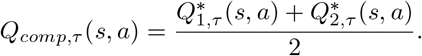

*The optimal Q-function* 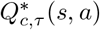 (*s, a*) *for the composed task is bounded by:*

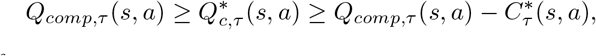

*where* 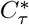 *is a fixed point of*

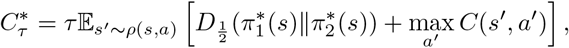

*where* 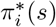 (*s*) *is the optimal Boltzmann policy for task i, and* 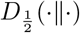 (·∥·) *is the Rényi divergence of order* 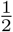.

## Notes

### Competing Interest Statement

The authors have declared no competing interest.

### Summary of Updates

The manuscript was revised to narrow its scope to the problem of composing multiple values. The previous results on the tail of the striatum have now been included in a separate manuscript dedicated to it.

